# System characterization of dynamic biological cultivations through improved data analysis

**DOI:** 10.1101/2021.05.14.442977

**Authors:** Gerben Roelandt Stouten, Sieze Douwenga, Carmen Hogendoorn, Robbert Kleerebezem

**Affiliations:** Department of Environmental Biotechnology, Delft University of Technology, Delft, The Netherlands; Systems Bioinformatics, AIMMS, VU University Amsterdam, Amsterdam, The Netherlands; Department of Microbiology, IWWR, Radboud University, Nijmegen, The Netherlands

**Keywords:** bioprocess characterization, soft-sensor, dynamic cultivations, PHA

## Abstract

Determining the functional development and dominant competitive strategy in microbial community enrichments is complicated by the extensive measurement campaigns required for off-line system analysis. This study demonstrates that detailed system characterization of aerobic pulse fed enrichments can be established using on-line measurements combined with automated data analysis. By incorporating the physicochemical processes in on-line data processing with a Particle Filter and kinetic process model, an accurate reconstruction of the dominant biological rates can be made. We hereby can differentiate between storage compound production and biomass growth in sequencing batch bioreactors. The method proposed allows for close monitoring of changes in functional behavior of long-running enrichment cultures, without the need for off-line samples, therewith enabling the identification of new insights in process dynamics with a minimal experimental effort. Even though a specific example application of the method proposed is described here, the approach can readily be extended to a wide range of dynamic experimental systems that can be characterized based on on-line measurements.

## Introduction

Enrichment studies are fundamental tools for understanding the microbial world. The earliest studies from Beijerinck and Winogradsky are based on restrictive growth media and gradients in space, which favor growth of particular microbes over others (1, 2). They were used to elucidate the role of microorganisms in biogeochemical cycles, where carbon, nitrogen, oxygen, phosphorus and sulfur compounds are converted in a wide range of redox reactions. Later enrichments established by Grant and Brock focused on the use of environmental factors to enrich microbes that flourish at high temperatures and salinities for example (3, 4). The countless combinations of selective media and environmental factors have unearthed a profound microbial richness (5). Although many microbes have been enriched in a wide range of conditions, all of these enrichments are based on imposing a specific selective pressure in either a continuous or batch system. Recent work has demonstrated that through imposing dynamic process conditions an additional wealth of microbial diversity is revealed, compared to traditional chemostat or batch cultivation (6–9). Essentially all natural environments are subject to ever changing conditions, from scarce nutrient availability to its sudden abundance and more regular day-night cycles leading to fluctuations in temperature and substrate supply. An abundant portion of microbes evolved under dynamic conditions and as such developed different functional strategies to exploit a competitive edge. For example, the alternating absence and presence of light, electron acceptor, carbon substrate, and growth nutrients provides a competitive advantage for specific microorganisms that depend on the production and consumption of storage polymers. We propose that a large part of the microbial diversity and available functionality is overlooked by studying microorganisms in absence of dynamics.

The exploration of dynamic cultivation systems increases the experimental complexity but gives more information on the development over time as each operational cycle can be analyzed and compared (10). The quantification of the variations between cycles requires data analysis which aims for identification of the time dependent stoichiometry and kinetics of the (biological) process in the bioreactor. The reconstruction of biological rates is complicated by two distinct dynamic properties. First, the change of environmental conditions, like cyclic nutrient pulses, result, by definition, in variable concentrations inside the bioreactor. The second dynamic is caused by the response time needed for measurable parameters to changes in the process, such as dissolved oxygen concentration or CO_2_ in the off-gas.

Throughout an operational cycle, microbes will respond to the changing environmental conditions and can exhibit different functional properties depending on the actual conditions. Capturing these changes in functional properties requires frequent measurements at short intervals. Combined with the fact that enrichment studies take weeks to months, manual sample analysis becomes both cumbersome and non-scalable. Online data from liquid probes and off-gas analysis is generally available but utilization of these measurements for on-line process identification beyond the observations of general trends is often limited, due to complicating physicochemical processes and measurements noise.

Here, we propose a method which allows high-definition system characterization, based solely on on-line measurements. This method combines a signal processing method (Particle Filter) and a physicochemical model (Figure 1). We demonstrate this method by reconstructing the respiration rates for a 100 day enrichment in an aerobic pulse fed bioreactor. The demonstration shows how the reconstructed respiration data can be used to ascribe microbial functionality to each of the 200 cycles. Herewith we aim to show the added worth of standardizing data reconsolidation in dynamic cultivations. As such, the approach suggested is widely usable in microbial cultivation, and can help us unlock new insights in metabolism, competition, and regulation of microorganism.

**Fig. 1.**
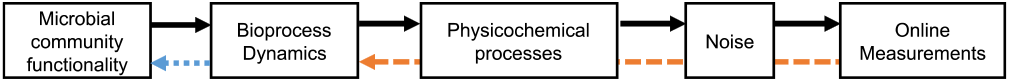
Schematic overview of the factors that influence on-line measurements. The goal of the Particle Filter (orange dashes) is to reconstruct the bioprocess rates as accurately as possible, taking into account the limitations resulting from system and measurement noise. Additional understanding of the metabolic processes is required (blue dots) to derive further biological information from the reconstructed bioprocess dynamics. In the example system, the reconstructed respiration rates are used to identify the dominant metabolic processes of the enrichment culture over time.

### Particle Filter and physicochemical model for respiration rate reconstruction

Mandenius and Gustavsson (11) reviewed several software sensor solutions that are used to derive new information from true sensors by combining them with software models. These models range from high mathematical signal processing complexity (12–14) to those with emphasis on biological system modeling that describe specific biological systems (15, 16). A general downside of these models is that either their implementation is of such complexity that they are difficult to realize and verify or that they are not suited for a general solution. Especially the mass transfer limitations that occur in systems with imposed dynamics (e.g.: pulse fed systems) and the difference in measurement frequency and accuracy of sensors lead to implementation difficulties (17, 18). Here we propose and demonstrate a model that is specifically suited for the reconstruction of respiration rates in dynamically operated bioreactors.

De Jonge and colleagues developed a physicochemical model that allows for the respiration rate reconstruction at per-second timescale of a chemostat culture perturbed by a single substrate pulse (19). Their model is meant as a research tool to facilitate state estimation and requires significant calibration, parameter estimation, and understanding of the Extended Kalman Filter utilized. More recent advancements in state estimation come in the form of Particle Filters where the implementation of the system physics are more accessible (20), and the results are more accurate than the Extended Kalman Filter, but the implementation is computationally more expensive (21). The Extended Kalman Filter and Particle Filter are used to estimate the internal states in dynamic systems based on partial observations in the presence of sensor noise. Both methods require a model of how the system changes in time as a response to changing inputs – in this case respiration rates. Here, the model incorporates the main physicochemical processes, which include inorganic carbon speciation, gas mixing in the bulk liquid and head space, delays and non-equilibrium phase transfer rates. While the Kalman Filter is the optimal signal processor for linear models, and it has been adapted to support non-linear systems in the form of the Extended Kalman Filter, its application requires local linearization, which increases the mathematical complexity during model development and model expansion and makes the model error-prone during strong non-linear behavior. The Particle Filter overcomes those issues by introducing many alternative starting points (particles) for each prediction step, this increases the computational complexity but reduces mathematical complexity, and allows adaption to non-linearities.

### Example system: Aerobic organic carbon degradation in a sequencing batch process

Microbial communities in pulse fed systems can exhibit at least two different metabolic functional strategies. Bacteria either grow on the pulsed substrate, or bacteria convert the pulsed substrate into an intracellular storage polymer. The latter type of hoarding bacteria can metabolize the stored polymers to catalytic biomass when the external substrate is depleted. Previous studies see a clear change in the community structure and functionality throughout the enrichment period of 30 to 100 days when pulse feeding is applied, resulting in a microbial community that is dominated by hoarding bacteria (23–26). The main intracellular compound that is produced in these studies is polyhydroxybutyrate (PHB), which is gaining significant interest because its material properties are similar to oil-derived plastics. These enrichment studies therefore predominantly focused on the final enriched community, reducing the interest in the development of the community functionality and structure throughout the enrichment. To allow more insights in the mechanisms behind microbial competition and succession in enrichment studies we aim to monitor and characterize the functional transitions of enrichment cultures in time based on on-line measurements. By reconstructing the respiration rates in the dynamic cultivation conditions important insight can be generated regarding the development of the dominant metabolic strategy.

## Materials and Methods

This section is divided in the materials and methods of (A) the parallel cultivation system that allows continuous on-line measurements and characterization of enrichment cultures, (B) the description and implementation of the Particle Filter and physicochemical model for the reconstruction of respiration rates based on on-line measurements, and (C) the example process of aerobic organic carbon degradation in a sequencing batch reactor on which we apply the proposed methodology.

### A. Cultivation setup for on-line characterization of enrichment cultures

In an effort to support high-resolution and reproducible enrichment studies a parallel bioreactor cultivation setup was designed and constructed (Figure S1). Considerable attention has been put in the flexibility of the hardware and software of the system to allow precise control over the cultivation conditions, which can be employed to impose a wide range of selective conditions on microbial cultivations.

The setup consists of eight identical 2.2L jacketed bioreactors (Applikon, the Netherlands), allowing temperature control through separate thermostats (RE630, Lauda, Germany). Each bioreactor is equipped with four feed/effluent pumps (WM120U, Watson Marlow, United Kingdom), and two acid/base pumps (MP8, DASGIP, Germany). Concentrated medium and water vessels are placed on balances (Mettler Toledo, The Netherlands). pH and dissolved oxygen probes for each reactor are connected to a measurement unit (PH8DO8, DASGIP, Germany). In-gas composition and gas flow for each individual bioreactor is delivered through a gas mixing manifold (MX44, DASGIP, Germany). Off-gas is led through a condenser, kept at 4°C through a cryostat (Lauda, Germany), and connected to the gas mass spectrometer (Prima BT, Thermo Fisher, USA). System operational pressure (typically 50-100 mbar overpressure) is controlled through a manually adjustable pressure control valve placed in the off-gas line. Gas recirculation of the headspace to the gas-inlet is facilitated by an adjustable membrane pump (Buerkle, Germany). Two six-blade Rushton impellers are stirred by on overhead drive, which is controlled by a stirring unit (SC4D, DASGIP, Germany).

A central control unit was custom made (HAL, TU Delft, The Netherlands) for analog control of the 32 feed/effluent pumps, and for the digital communication with the DASGIP units, balances, thermostats and mass spectrometer. Custom logging and scheduling software (D2I) allowed for fine control over the dosing mechanism, gas profile switching, incorporation of off-gas and off-line measurements as system event triggers, and general automatization of system variations and calibrations.

#### Mass spectrometer tuning

Gas composition going in and coming out of the reactors was analyzed with on-line mass spectrometry (Prima BT, Thermo Fisher, USA). The fast switching of the mass spectrometer allowed for frequent gas composition monitoring, where the in-gas and the off-gas of eight reactors are measured every 3 minutes. The Prima BT mass spectrometer contains a small flow cell which is continuously analyzed at different m/z values. The composition of calibration gasses can be analyzed to ppm level because the gas composition remains constant. During dynamic bioreactor operation the off-gas composition changes continuously. As a result, the gas composition in the flow cell changes throughout the measurement window of the mass spectrometer. Therefore, a tradeoff has to be made between analysis time and accuracy. In the current configuration we use 10 seconds of stream flushing and 8 seconds of measurement per channel (2 seconds per m/z: 28, 32, 40, 44). This results in the expected errors in the gas composition measurements between 10-300 ppm depending on the gas compound mole fraction. With eight gas_OUT_ and one gas_IN_ channel, the total time between measurements for each system is less than three minutes.

#### Online data processing pipeline

Online data is processed by filtering out erroneous measurements collected during operational disturbances (e.g.: bioreactor cleaning and maintenance), and correcting for the delayed off-gas composition measurement due to the off-gas plug-flow from the bioreactor to the mass spectrometer. Generally, filtering data and constructing datasets from raw measurement data is a manual task, due to many ambiguities that arise from differences in experimental setup and operation. Given that the parallel cultivation setup developed in this work spans eight analogue bioreactor setups that run highly comparable cultivations, this task could be largely automated. By processing the raw on-line measurement with a heuristic based, data filtering, and data pruning approach the task of data processing was significantly accelerated. Python3 code (filteredcycle.py) is included in the on-line code repository at: https://github.com/GRS-TUD/d2i, with examples of common disturbances, and their heuristics. From these filtered datasets the respiration rates (oxygen uptake rate and carbon evolution rate) are reconstructed throughout each cycle using the Particle Filter with the physicochemical model.

### B. Particle Filter and physicochemical model for reconstructing Bioprocess Dynamics

#### Reconstructing the respiration rates with a Particle Filter

The Particle Filter is a 3-step, prediction-update-resample, state estimation method that uses *N* samples from a distribution around the current (k) best estimate of the system state to predict the changes to the system state through an underlying model (detailed implementation in Supplementary Material 2). With each calculation cycle a step (k+1) in time is made. Each particle represents a possible oxygen and carbon dioxide respiration rate (hidden state) which are used as input in the physicochemical model to predict the new compound concentrations and measurements. During the update step the uncertainty of *N* new state predictions and the uncertainties of the actual measurements (observation) are taken into account to make a weighted distribution of the most likely system state using Bayes’ theorem. As *N* tends to infinity this method converges to a system state as expressed by a probability density function. During the resample step, new particles are chosen to prevent highly unlikely system states (particles) to take-up calculation time and to increase the resolution surrounding the current best state estimates. This re-sampling step requires a resampling algorithm, here we chose to implement the systematic resampling algorithm from Douc et al. (27).

The Particle Filter requires two additional inputs, (i) the covariance matrix of the process represents how fast the biological respiration rates can change, and (ii), the covariance matrix of the measurement captures correlations and noise of measurements. These matrices should be derived for each specific setup and experiment.

#### Determining the Particle Filter covariance matrices and initial state

The aim of the Particle Filter is to reconstruct the respiration rates of the biological community. During the prediction step each particle represents the rate of change of the current respiration uptake rates, 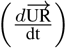,where the values are normally distributed around 0, and with a specific variance for both the oxygen uptake rate (OUR) and carbon dioxide uptake rate (CUR). Negative uptake rates equal positive production rates. The process variance for the OUR and CUR allows the model to emulate the development of the respiration rates and is chosen in such a way that the observed increase and decrease of the respiration rates on a substrate pulse can be achieved. If the values are too small, the model cannot respond quickly enough to changes in respiration rates, and if the values are too large a significant fraction of the particles will take on values that diverge far from the true mean.

In most aerobic biological systems a strong correlation exists between oxygen uptake and carbon dioxide production, incorporating this knowledge reduces the computational effort because fewer particles are required to reconstruct the respiration rates. To show the general applicability of the model, the results shown here assume no additional knowledge regarding the respiration covariance. Therefore, the initial estimates and covariance matrix are diagonal matrices. The modelled biological system does not involve nitrogen fixation or production (NUR = 0). The process covariance matrix (mol^2^ s^-2^) for the system modelled here is given by:

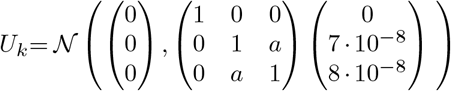

With U_k_ representing the covariance matrix for (nitrogen, oxygen and carbon dioxide uptake rates), where *a* is 0 if no correlation between OUR and CUR is assumed, and *a* is −1 assumes a perfect correlation.

All analyses contain some measurement uncertainty. This uncertainty is taken into account to make a weighted average between the predicted and measured values for the system state. The measurement noise covariance for the off-gas (N_2_, O_2_, CO_2_) measurements (expressed as ppm^2^) is based on the accuracy of the mass spectrometer:

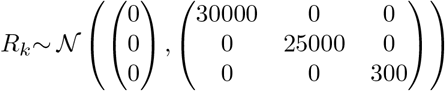

With R_k_ representing the covariance of head space gas mole fraction (N_2_(g), O_2_(g), CO_2_(g)).

The initial state of the system (gas concentrations and dissolved carbon species at t=0) is important for model predictions. The Particle Filter allows the initiation of N different initial states which converge to the most probable state. Simulation with the Particle Filter were performed with 10.000 particles, initially uniformly distributed between 0 and 10^−6^ mol_O2_ s^-1^, and between −10^−6^ and 0 mol_CO2_ s^-1^.

#### Physicochemical processes for gas exchange

The mole fraction of gas species in the headspace (h) and in the bubbles (b) can be described in vector notation (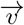 with ⊙ as element wise multiplication operator). The change in the concentration of the dissolved gas species in the broth depends on the transfer to and from the gas phase 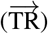 and the uptake or production by micro-organisms 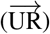.

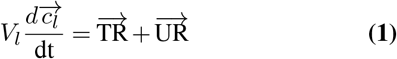

The transfer rates 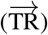 depend on the partial pressure of the gasses in the bubbles

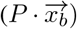, the concentration in the liquid 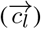 and the mass transfer coefficients 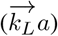.

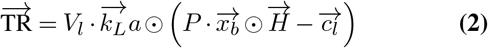

The transfer of gas to or from the bubbles changes the molar gas flow rate from the bubble to the headspace 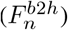. It may practically be assumed that all gas in the reactor is saturated with water vapor (P_W_). Headspace recycling 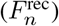 bypasses the off-gas condenser to appropriate the drying capacity to off-gas flowing to the mass spectrometer (Figure 4).

**Fig. 2.**
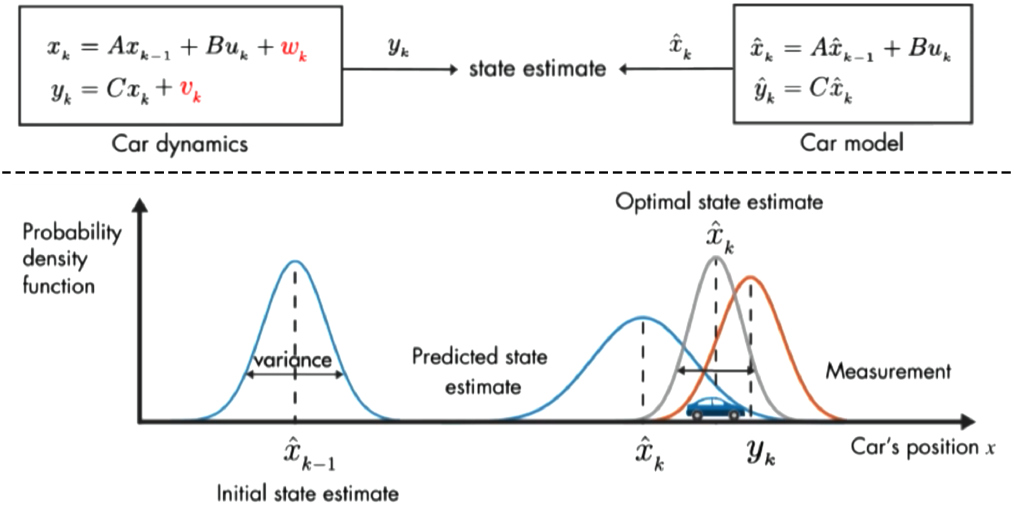
Schematic overview of the Kalman Filter when applied to tracking a car. By making a model of how a car behaves with respect to the laws of physics we predict where it will end up in the next time step. This prediction has some uncertainty (blue bell shape), which increases with increasing time between steps. When measurements are available, we take their uncertainty into account (red bell shape). The best estimate for the car state is than the statistical overlap of the two probabilities. **To complete the analogy, the problem that we are solving in our model is trying to get a good estimate of when and how far the gas paddle and breaks are pressed based only on GPS data**. Image adapted from *Understanding Kalman Filters* (22)

**Fig. 3.**
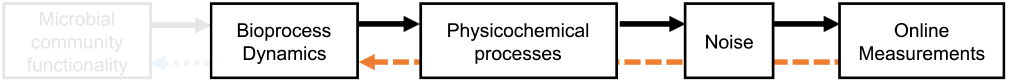
The goal of the Particle Filter (orange dashes) is to reconstruct the bioprocess rates as accurately as possible, taking into account the limitations resulting from system and measurement noise.

**Fig. 4.**
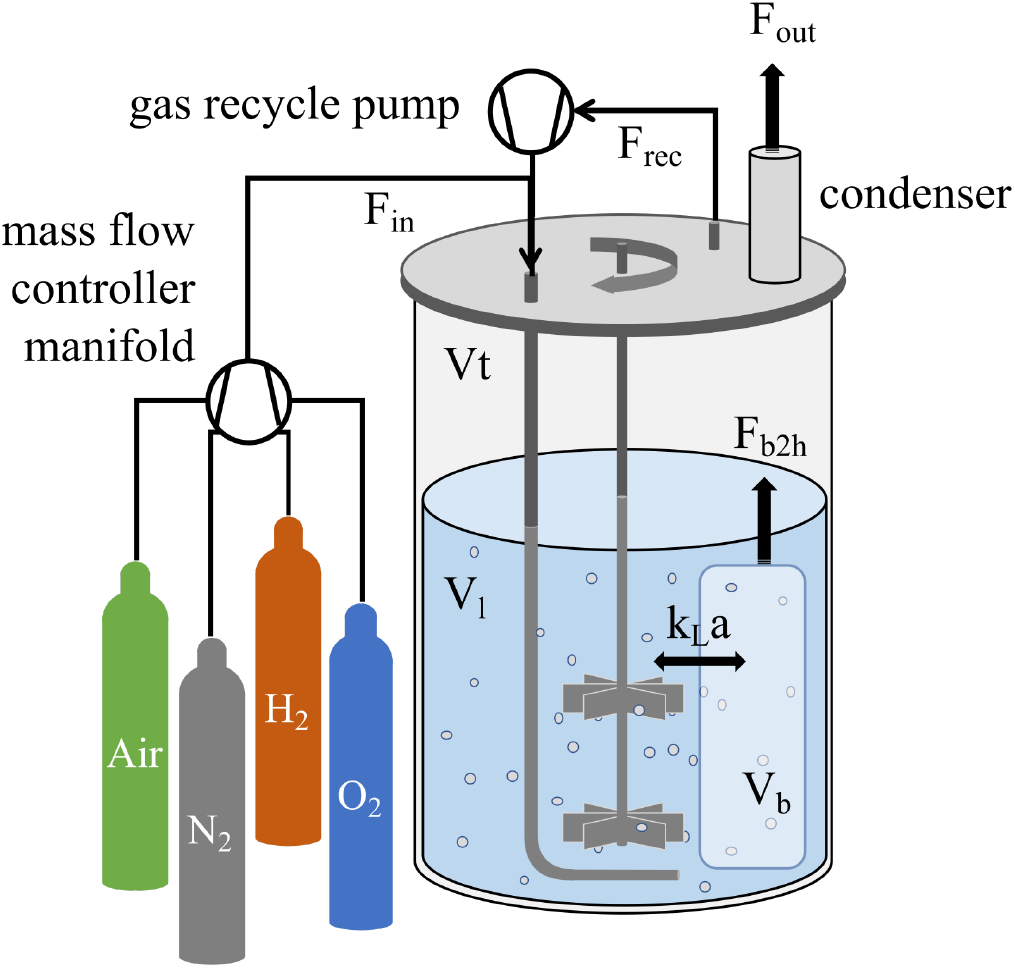
Schematic overview of the gas holdup and gas flow in a bioreactor. Dry gas (F_in_) from the gas mixing manifold combines with wet gas from the headspace (F_rec_), and is led into the bioreactor. The bubbles take up a specific volume (V_b_) in the liquid (V_l_), gas exchange takes place between the gas bubbles and the liquid through rate parameter (k_L_a). Gas in the bubbles saturates with water and a net flow of wet gas (F_b2h_) takes place from the bubbles to the headspace (V_h_). Wet gas is dried in the condenser and flows to the mass spectrometer (F_out_).

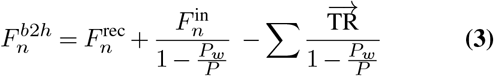

The change in the composition of the gas in the bubbles depends on the gas composition flowing in (combining new ingas with potential headspace recirculation).

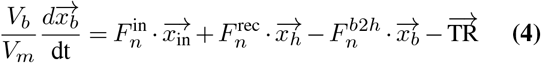

The change in the composition of gas in the head space depends on the molar gas flow rate and the composition of the bubble and headspace gasses.

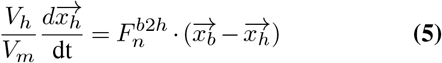

The equations above describe the change in concentrations and mole fractions of gasses in the liquid and gas phase. For carbon dioxide an additional factor needs to be taken into account, the speciation of inorganic carbon as CO_2_(aq), H_2_CO_3_, HCO_3_^-^, and CO_3_^2-^ (Figure 5). Speciation depends on forward and backward reaction rate constants (lowercase k) and equilibrium constants (uppercase K), their values have been adapted from the comprehensive studies by Harned and colleagues (28, 29), and Wang and colleagues (30)^1^.

**Fig. 5.**
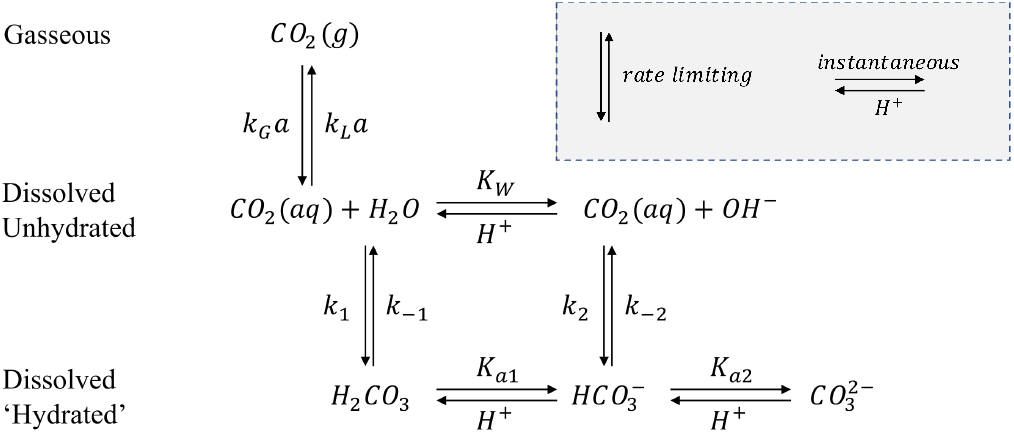
Reaction scheme of inorganic carbon in aqueous solution. Reactions that take place over the vertical axis are rate limiting, and reactions taking place over the horizontal axis are instantaneous protonation equilibria. In the gas-liquid exchange the k_G_a and k_L_a have the same physical meaning but are written from the perspective of the gas or liquid phase.

The protonation equilibria are considered instantaneous with respect to the timescales of our experiments (31). The hydrated carbonate species (carbonic acid, bicarbonate, and carbonate) can therefore all be expressed as a fraction (f_i_) of the total hydrated inorganic carbon 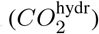, and depend on the pH.

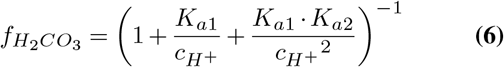

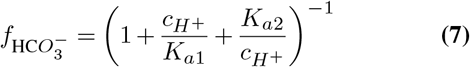

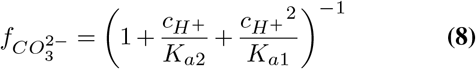

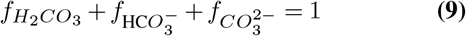

The kinetics of inorganic carbon speciation can be seen as the net transfer of carbon between the hydrated (H_2_CO_3_ and HCO_3_^-^) and dehydrated (CO_2_(aq)) species.

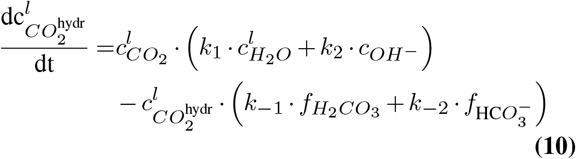

The three processes that therefore contribute to the dissolved CO_2_ concentration in the reactor broth are: the carbon dioxide evolution from microbial activity, the carbon dioxide transfer between the gas and liquid phase, and the (de)hydration reactions for carbon dioxide speciation. The initial two processes are already captured in equation (1), and therefore the (de)hydration rate has a reducing (-=) effect on the concentration of dissolved CO_2_.

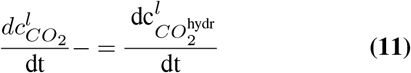

Furthermore, the broth in a bioreactor is a non-ideal solution, therefore compound activity needs to be used instead of concentrations for charged species. The extended Debye-Hückel equation is used as approximation for the activity coefficients for solutions with an ionic strength (I) lower than 100 mmol L^-1^:

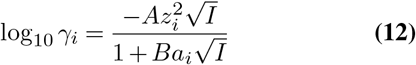

Here, *γ*_i_ represents the activity coefficient and z_i_ is the charge of ion *i* (H^+^, OH^-^, HCO_3_^-^, CO_3_^2-^). A, and B are temperature dependent constants, and have values 0.496 (kg^0.5^ mol^-0.5^), and 0.325 (Å^-1^ kg^0.5^ mol^-0.5^) at 30 °C. The value for *a*_*i*_ is the effective diameter of the ions (H^+^ 9Å, OH^-^ 3.5Å, HCO_3_^-^ 4Å, CO_3_^2-^ 4.5Å (32)). The ionic strength of the medium used in the example (see section Material and Methods C) is approximately 75 mmol L^-1^, bringing the activity coefficients for protons, hydroxyl, bicarbonate, and carbonate to 0.84, 0.79, 0.8, and 0.41, respectively.

### C. Application of proposed methodology on enrichment of pulse fed aerobic sequencing batch bioreactor (example)

#### Sequencing batch bioreactor

Throughout this manuscript the experimental data of a single enrichment study is used as example. The bioreactor was operated aerobically, for 200 cycles (100 days), with acetate as sole carbon source and electron donor (40 mCmol pulse per cycle). The working volume of the bioreactor was 1.4 L, and every 12 hours half of the bioreactor content was discharged and replaced by 0.7 L growth medium (7.1 mM NH_4_Cl, 0.7 mM KH_2_PO_4_, 0.2 mM MgSO_4_.H_2_O, 0.2 mM KCl, and 1.5 mL L^-1^ trace solution (33)). pH was maintained at 7 ± 0.1 by use of 1 M HCl, and 0.5 M NaOH. Gas flow in the reactor was compressed air at a flow rate of 400 ± 2 mL min^-1^. This enrichment was part of a study that investigated the influence of temperature on the development and performance of PHB enrichments by operating eight bioreactors in parallel (34).

#### Cycle measurements

Cycle measurements of SBR operations were performed to fully characterize the processes occurring in the system and verify system characterization efforts based on on-line data. Measurements are performed as described by (35). Samples taken from the reactor for analysis of acetate and ammonium were immediately centrifuged and filtered with a 0.22 µm pore size filter (PVDF membrane, Millipore, Ireland). PHB and biomass dry weight measurements are performed on 15 mL bioreactor samples, after centrifuging at 5000 g and freeze-drying of the pellet.

#### Key indicators for characterization of functional system behavior

In a pulse fed bioreactor, the extracellular carbon will be depleted at a certain time, this carbon period is defined as the feast-phase. The period in which no extracellular carbon is present is defined as the famine-phase. Three functional characteristic key indicators were derived that summarize the functional behavior in an aerobic pulse-fed sequencing batch bioreactor (Figure 8). All key indicators can be derived from the reconstructed respiration rates.

**Fig. 6.**
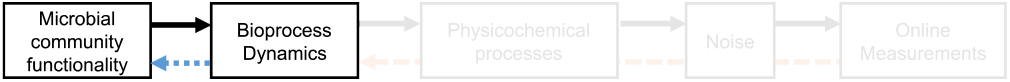
The Particle Filter reconstructs bioprocess dynamics in the form of respiration rates. These rates can be used to further characterize the microbial community functionality when additional information about the system is considered (blue arrow). This process is system specific, and here we describe how the Particle Filter can be used for the pulse fed aerobic bioreactor systems.

**Fig. 7.**
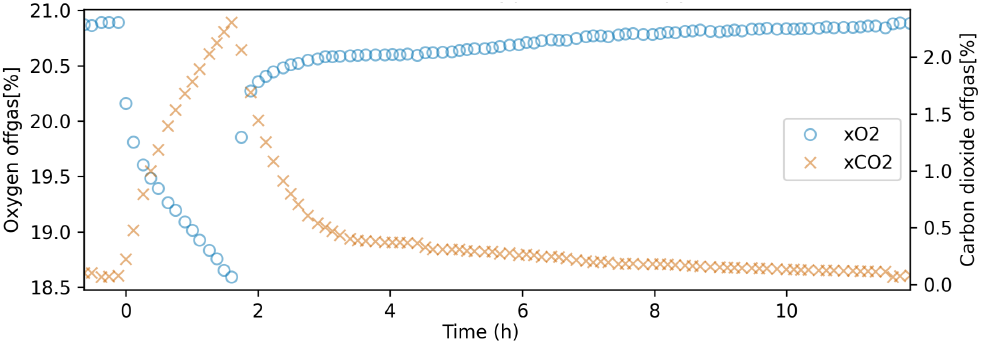
Example off-gas measurement of a single operational cycle. At t=0 a substrate pulse is given, which around t=2 h is completely consumed leading to a drop in respiration activity. The goal of the Particle Filter is to convert these measurements to the biological respiration rates.

**Fig. 8.**
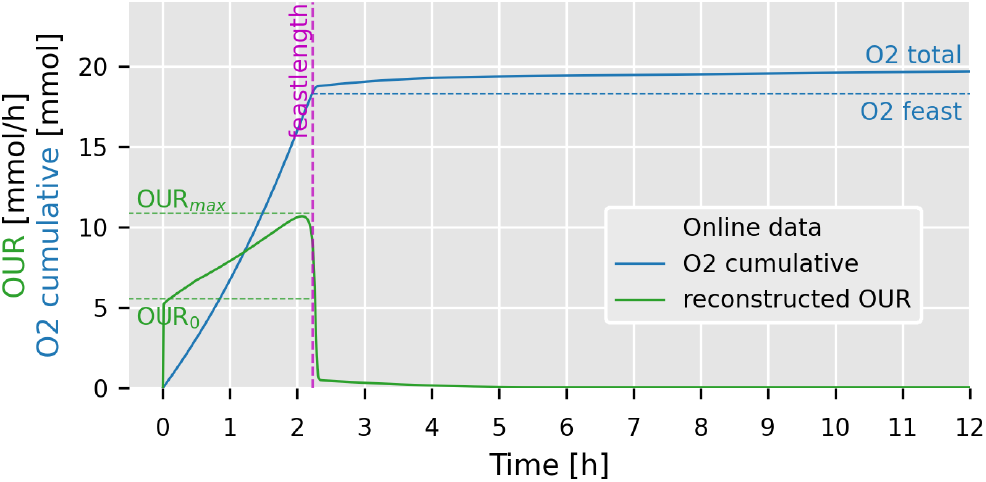
Schematic representation of how the reconstructed oxygen uptake rate (OUR in green) can be used to derive cycle specific key indicators. After the substrate pulse at t=0, a change in OUR is observed until the substrate is depleted at the end of the feast phase (feastlength in purple). By integrating the OUR over the cycle (O2 cumulative in blue), it is possible to estimate the fraction of total oxygen that is consumed during the feast phase.

- The *feast-length* is derived as the inflection points of the OUR, CUR and dissolved oxygen (DO). By reducing the number of points in the respiration curves with the Ramer-Douglas-Peucker algorithm (*ε* = 0.01) the curve is simplified (36). From this curve the slope was calculated for each segment, and the end of the feast-phase is positioned at the intersect of the most aggressive slope and the first prior segment with opposite sign. Offline measurement of acetate verified that this method closely predicts the true end of feast (Supplementary Material 4).
- The *O*_*2*_*feast%* is derived as the fraction of oxygen that is consumed in the feast-phase over the total oxygen consumed in the cycle. This value follows from the feast-length and OUR data, and is an indicator for the amount of respiration that takes place in the feast versus the famine phase. Where a lower *O*_*2*_*feast%* indicates that significant respiration takes place in the famine phase, which is assumed to be due to respiring storage compounds.
- The *OUR* _*ratio*_ is derived as the ratio between the oxygen uptake rate at the beginning of the feast-phase (OUR_0_) to the maximum observed oxygen uptake rate (OUR_max_). A strong increase in the respiration activity throughout the feast phase indicates that the catalytic capacity of the system is increasing, which is likely caused by growth. While, if the respiration activity remains constant throughout the feast phase, very limited growth can be expected and therefore the majority of the substrate is likely directed to storage polymers.

#### Identification of non-gaseous compounds

The Particle Filter is used to reconstruct the respiration rates of a biological system. These respiration rates reflect the changes in biological activity of the biomass in the system, but they represent only a limited number of compounds in the biological conversions of the system. To utilize the reconstructed respiration rates further, additional information on the cultivation and expected conversions is required. Here we will show, for this specific case, how the data can be used to reconstruct the dominant compound profiles in a sequencing batch bioreactor that is pulse fed with acetate.

The respiration profiles are used to perform carbon and electron recovery calculations throughout each cycle. The following assumptions are made with respect to the carbon and electron utilization. (i) Acetate is the only carbon and electron donor. (ii) All acetate is converted to either catalytic biomass (CH_1.8_O_0.5_N_0.2_), storage polymers (PHB: *n* x C_4_H_6_O_2_), or carbon dioxide. (iii) Storage polymers are transient, and are therefore consumed by the end of the cycle. (iv) Oxygen is the only electron acceptor. (v) The system operates in a pseudo steady state. (vi) PHB and biomass yields are taken from Johnson et al. (35). Furthermore, the systems are only carbon substrate limited, pulsed with acetate at the start of the cycle (t=0h), and 50% of the reactor content is exchanged with new medium at the end of the cycle.

#### Biomass and PHB production during the feast phase

An approximation of the amounts of catalytic biomass and storage polymers that are produced during the feast phase can be made, based on the oxygen consumption and carbon dioxide production during the feast phase. The three biological processes involving acetate uptake during the feast phase are: biomass production, storage polymer production, and cellular respiration. The stoichiometries are taken from Johnson et al. (35) and given below in carbon mole ratios [C-mol].

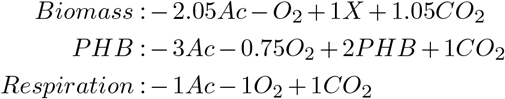

By combining these stoichiometries in the electron and carbon balance we can derive an analytical expression for the amounts of biomass and PHB that are produced during the feast phase. Simplified derivations are shown below, where the contribution of respiration is neglected during the short feast phase.

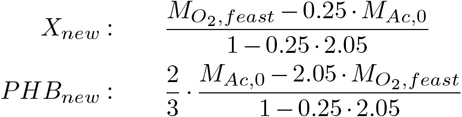

These equations demonstrate that the oxygen consumed in the feast phase can be used to give a direct insight in the prevalence of storage polymer production for each cycle. Analogues equations can be derived for total carbon dioxide. Here, the change in the respiration rate (e.g. OUR profile) is not yet utilized, this is important because different profiles can yield similar total respirations. Therefore, more insights in biological functionalities could be derived by reconstructing the full compound profiles.

#### Compound profiles

To reconstruct the compound profiles throughout the whole cycle we can make use of the biological process model of Johnson et al. (35). This model takes as input off-line measurements of acetate, ammonium, biomass, and PHB, and combines them with on-line oxygen and carbon dioxide measurements. That model focuses on data reconciliation to derive kinetic parameters. We use the same model, but solely based on the on-line measurements to determine the acetate, ammonium, biomass, and PHB profiles. The frequency and resolution of our measurements do not allow for the determination of affinity constants, but do allow for the determination of dominant compound profiles. Additionally, it serves as a verification tool for off-line measurements.

The model requires initial estimates for the compound amounts, which are derived from the respiration measurements when combined with the principle assumptions described above. The initial amount of biomass in carbon-mol (M_X,0_) can be estimated from the total carbon dioxide production, the dosed acetate (M_Ac,dosed_), and applied exchange ratio (f_ER_). This also gives an estimate for the initial amount of ammonium, by assuming a fixed catalytic biomass composition (Y_NX_ = 0.2 N-mol / C-mol), and known amount of dosed ammonium (M_NH4,dosed_).

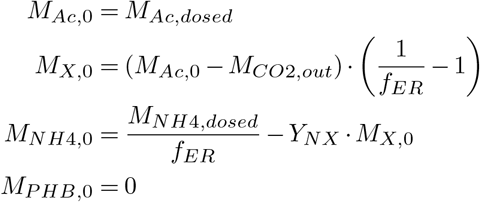

With these initial estimates and the respiration profiles the compound profiles are constructed from which estimates for biomass specific rates follow in the kinetic and stoichiometric PHB model of Johnson et al. (35), but now solely based on on-line measurements.

During the liquid exchange step, 50% of the biomass is removed from the bioreactor. Therefore, it can be expected that the increase in respiration activity during the feast phase (OUR_ratio_ should minimally be 0.5. Lower ratios indicate that activity is lost during the cycle. This decrease can be modeled by allowing a distinction in active and inactive catalytic biomass, where the fraction at the beginning of the cycle is given by.

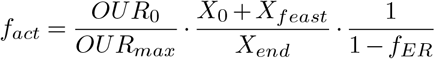

## Results

A microbial enrichment was established from activated sludge in an aerobic sequencing batch reactor, characterized by the supply of the limiting carbon substrate (acetate) at the beginning of every 12-hour cycle (34). Over a period of 200 cycles the functional performance of the bioreactor was characterized by on-line measurements. High frequency off-gas measurements were used to reconstruct the biological respiration rates through a physicochemical model and a particle filter. Reconstructions of two cycles with distinct functional performance are shown in Figure 9.

**Fig. 9.**
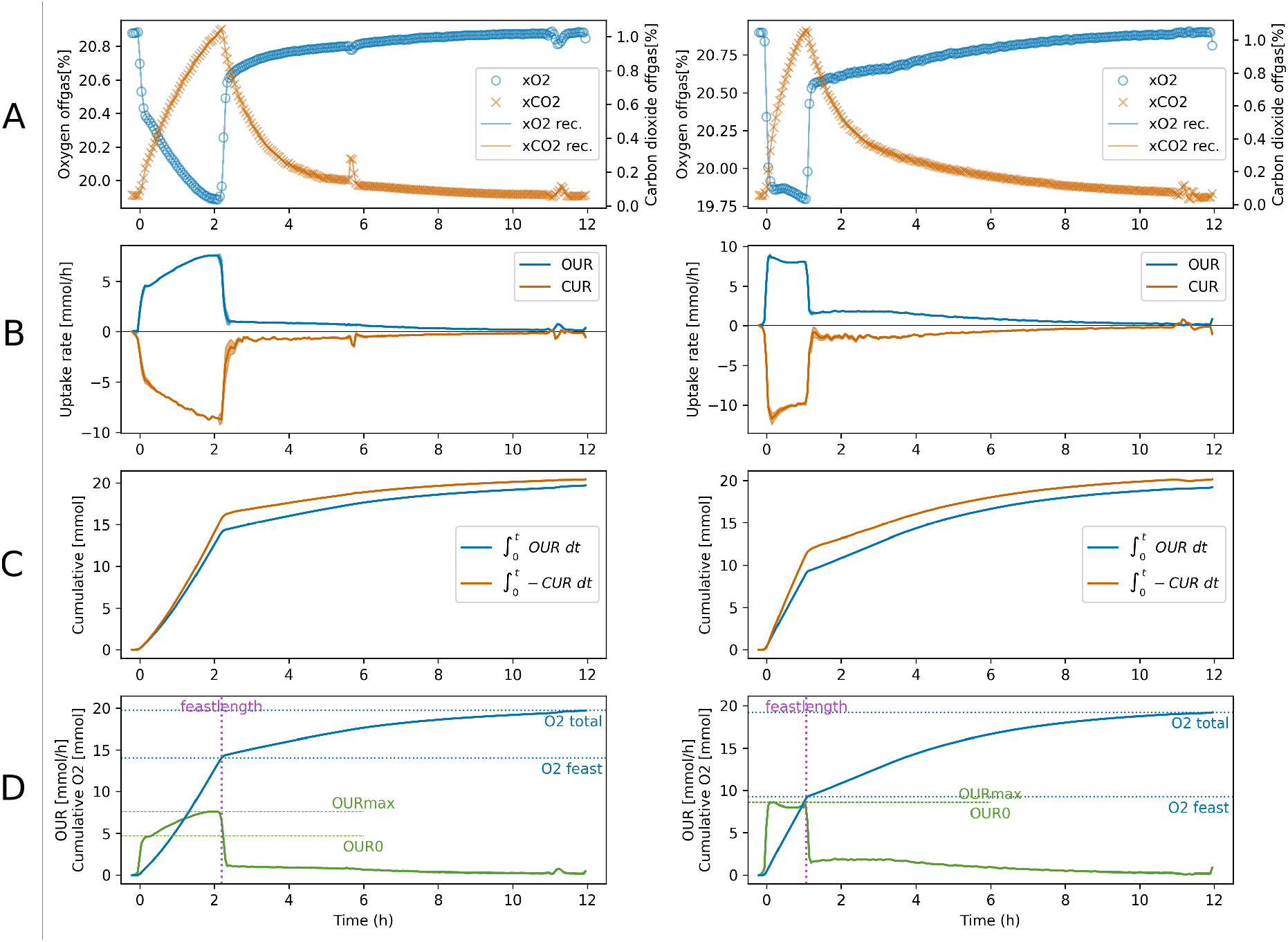
Reconstruction of microbial respiration rates from on-line measurements in a pulse-fed sequencing batch bioreactor exhibiting predominant growth and partial storage polymer production (left), or predominant storage polymer production (right) as response to the acetate pulse at t = 0 h. Sub-figures A show the off-gas measurements as symbols and the reconstructed off-gas profiles as continuous lines. Sub-figures B show the reconstructed respiration rates, where the line-widths represent the 95% confidence interval. Sub-figures C show the cumulative oxygen consumption and carbon dioxide production. Sub-figures D show the key indicators that were derived from the on-line data.

### Functionality development over time

By processing the on-line data of 200 cycles and calculating the key functionality indicators for each cycle (sub-figures D in Figure 9) it becomes possible to observe shifts in functionality over time (Figure 10). The observed shifts were verified with off-line measurements of PHB, microscopy, and microbial community composition through 16S rRNA gene sequencing. For an in depth analysis of the correlation between functionality and community developments and how they are influenced by temperature see Stouten et al. (34).

**Fig. 10.**
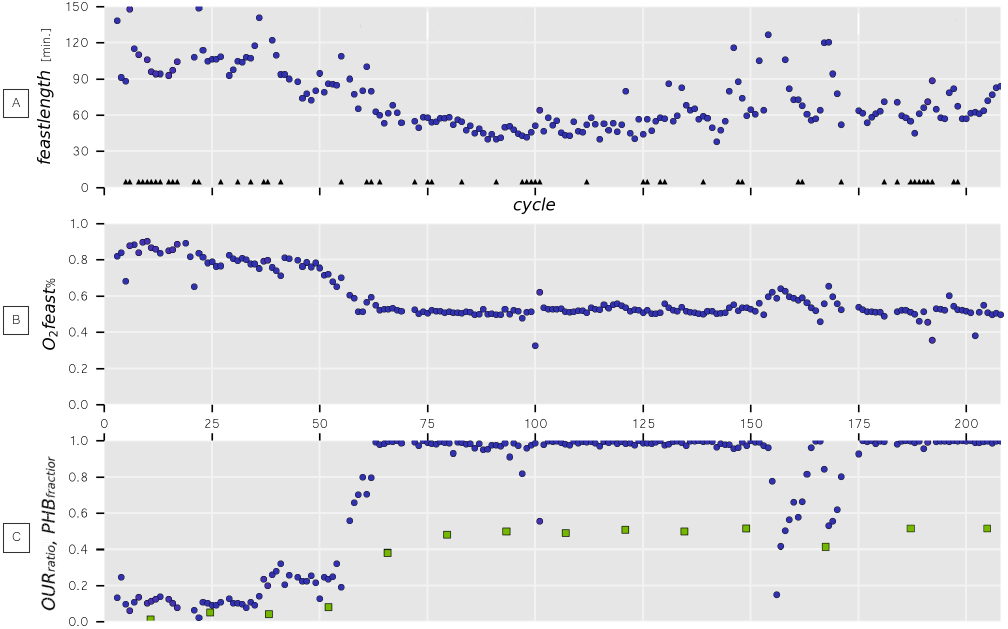
Overview of key functionality indicators over 208 enrichment cycles. Characteristic values (feast length, O_2_ feast%, and OUR_ratio_). Also shown is the intermittently off-line measured PHB fraction (g_PHB_/g_VSS_) at the end of the feast phase (solid green squares). Clearly visible are the functional transition between cycle 50 and 62, the upsets between cycle 150 and 175, and the overall stability of the enrichment.

### System characterization: compound profiles

The reconstructed respiration rates, combined with key indicators (Figure 9) are used as input in the kinetic and stoichiometric biological model of Johnson et al. (35) to approximate the compound profiles of acetate, biomass, PHB, and ammonium. In Figure 11 the reconstructed profiles are shown and compared with off-line measurements. The reconstructed compound profiles during the feast phase closely match the off-line measurements. During the famine phase a distinct difference is shown for the predicted PHB and biomass profiles and measurements. The final modeled biomass at the end of the cycle does match the measurements.

**Fig. 11.**
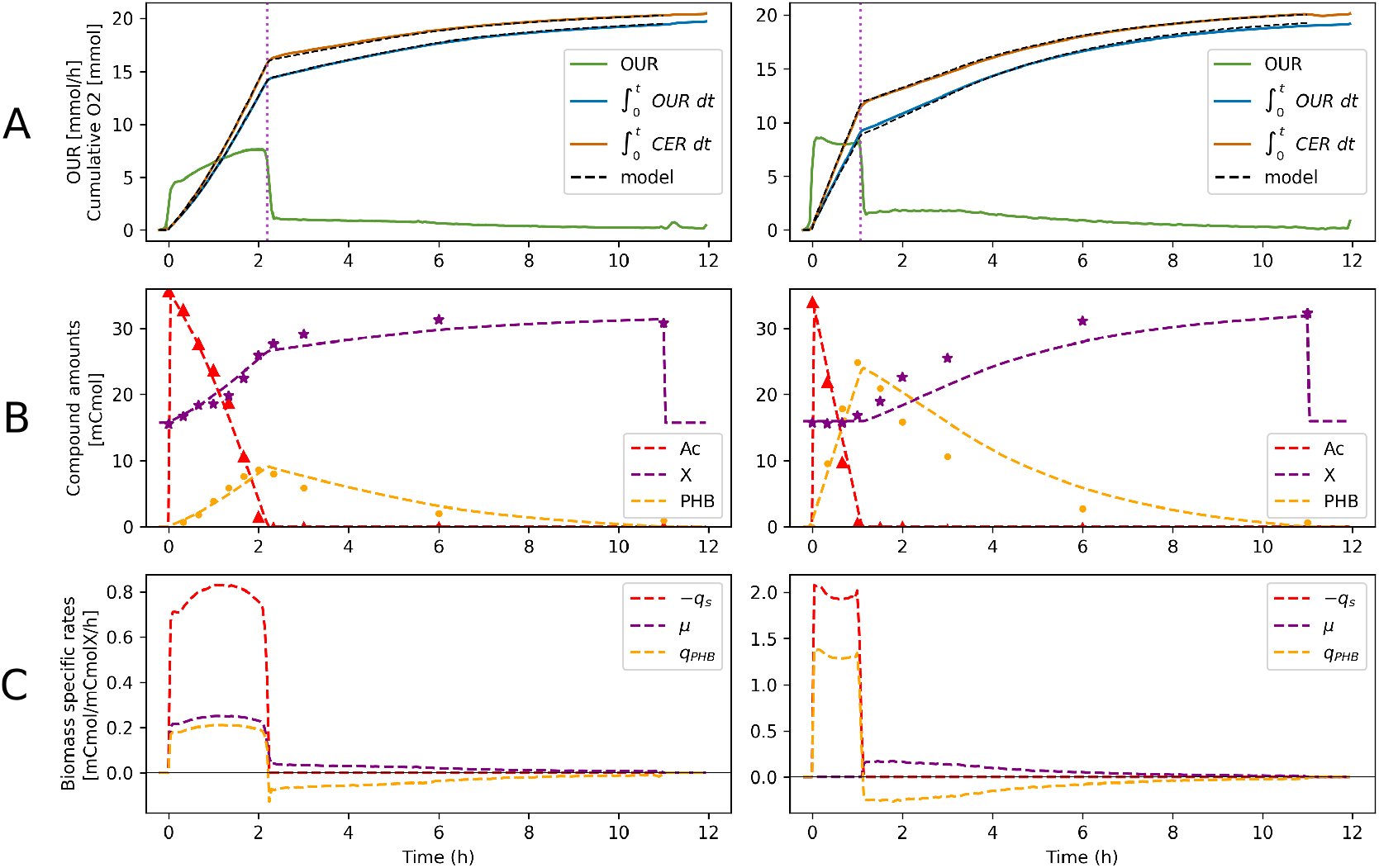
Reconstruction of compound profiles based on on-line data for two cycles during the enrichment of PHB producing microbial communities. The shown cycles are those from Figure 9, where the left system reaches a maximum PHB content of 25 wt%, and the right reaches a maximum PHB content of 55 wt%. Sub-figures A show the reconstructed oxygen uptake rate, cumulative oxygen consumption, and carbon dioxide production. The dashed lines overlapping the cumulative data indicate the modeled compound profiles for oxygen and carbon dioxide. Sub-figures B show the modeled compound profiles for acetate, biomass, and PHB - all based solely on on-line measurements. Also shown are off-line measurements as symbols. Sub-figures C show the on-line characterized biological rates for acetate uptake (*q_s_*), biomass production (*µ*), and PHB production and consumption (*q*_PHB_).

With the stoichiometric growth reaction on PHB present in the model, the PHB consumption profile from the off-line measurements cannot be achieved without a significant deviation from the reconstructed respiration profiles. In general, the model is able to clearly represent the compound profiles, and thereby also gives insights in biological rates. It should be noted that the biological rates as represented by the model (Figure 11C) reflect all biomass as if the community is composed of a single species that exhibits the same behavior.

## Discussion

### General applicability

The main objective of this study was to develop a parallel cultivation platform and the analytical tools to facilitate data processing during research with dynamically operated bioreactors. The eight identical bioreactor setups were equipped with a wide range of controls to allow mimicking and imposing exact environmental selection pressures. High-resolution off-gas measurements through mass spectrometry allow for non interfering monitoring of system dynamics. The off-gas signal is a convolution of the bioprocess gas uptake rates and several physicochemical processes. The developed Particle Filter model proved valuable in the reconstruction of actual respiration rates based solely on on-line off-gas measurements (Figure 9). The purpose of the Particle Filter is demonstrated by reconstructing the respiration rates in an aerobic, pulse-fed bioreactor throughout a cultivation period of 100 days. The resulting reconstruction of the respiration rates made it possible to identify moments where the dominant metabolic functionality of the microbial community changed (Figure 10). These characterizations were used to correlate changes in community composition with changes in observed bioreactor performance (34). The Particle Filter implementation demonstrated here, was used to reconstruct the respiration rates in a wide range of dynamically operated bioreactors, varying from aerobic pulse-fed bioreactors (34), anaerobic bubble columns (37), anoxic N_2_O respiration (38), aerobic/anaerobic fermentations (39), isotope labeled studies for identification of novel pathways (40), and in chain elongation studies (41). The applicability of this reconstruction tool contributes to the identification of functional properties of biological processes in dynamic environments. It furthermore enables the identification of process instabilities (Figure 10) which are typically associated with operational irregularities, but in this case can directly be attributed to biological dynamics.

In future work, the tool can be used for improved experimental design of dynamically operated bioreactors, including periodic disturbance experiments, design of real-time system response as process triggers, and eventually enables improved process model calibration and identification of parameter values and their sensitivity.

### Performance of the suggested approach in pulse-fed aerobic bioreactor system

In the pulse-fed aerobic bioreactor system investigated in this work three performance indicators (feastlength, O2_feast_%, and OUR_ratio_) were identified that allow for monitoring of the system development in time. These identifications were based on minimal knowledge of the biological processes in the system. No process yields, nor reaction stoichiometries were required to identify periods where transitions in the dominant metabolic functionality occurred. The validity of the automated on-line performance indicators was assessed by manual off-line cycle measurements, where samples from the reactor broth were analyzed for biomass, acetate, ammonium, and PHB contents. The data from these cycle measurements shows high agreement with the identified respiration rates (Figure 11). These observed relationships allowed the indirect assessment of PHB productivity throughout the enrichment (Figure 10), solely based on on-line data.

For the pulse-fed system described here, a kinetic process model is available that determines affinity constants and biomass specific rates for acetate uptake, PHB production, biomass growth, and maintenance (35). That model predominantly utilizes data collected off-line during cycle measurements, data from on-line off-gas measurements are assigned a low contributing weight (<10% relative to liquid measurements) during parameter fitting. This could be explained by the significant difficulty of incorporating on-line data of dynamic processes. Their less detailed physicochemical model of the bioreactor make it difficult to consolidate the on-line and off-line measurements, therefore the off-gas measurements were not in agreement with the modeled off-gas values. Furthermore, their kinetic process model describes the observed processes by a single set of stoichiometric and kinetic parameters. With the improved confidence in the reconstructed respiration data from the Particle Filter, new insights in the microbial functionality could be achieved. For example, the results in Figure 11 suggest that the biomass growth on PHB during the famine phase consists of at least two distinct processes, i.e.: the uptake of ammonium, and the cellular growth, including division. This distinction of PHB as carbon source (building block) and as energy source is shown in Figure 11, where the PHB consumption rate based on respiration data is lower than what is measured off-line. If both measurements are to be trusted, less PHB is respired in the early stage of the famine phase than is assumed to be necessary for growth, as described by the stoichiometries of the model. This increased confidence in reconstructed respiration profiles can thereby assist in our understanding of biological systems.

Off-line sampling strategies for the monitoring of enrichment systems and development of community characteristics over time are exceptionally laborious, even more so for dynamically operated systems. The Particle Filter combined with a process model allows for high resolution system characterization, but with a reduced parameter estimation compared to off-line measurements. The Particle Filter can be utilized for any biological system where the respiration data reflects the biological processes. In future work, the Particle Filter may be extended with detailed process models involving different microbial functionalities, and variable abundance of microorganisms, thereby attempting to model microbial ecosystem development, which in turn can be linked with molecular data on the development of the microbial community in the process (34).

### Limitations and extensions of the Particle filter and physicochemical model

The Particle Filter will result in respiration profiles that best fit the measured off-gas concentrations and the dynamics as imposed by the process variance. Without any additional knowledge of the system this can result in periods where biologically unlikely uptake rates are modeled, for example carbon dioxide fixation and oxygen production in heterotrophic systems. Such behavior shows up in the Particle Filter when fast changes of the system are not covered by the process variance and number of particles, for example at the exact moment of the substrate pulse, or at the depletion of substrate. This behavior requires an adjustment to the process variance parameter (*U*_*k*_). The predication uncertainty increases when the underlying biological process changes faster than is covered by the process variance and off-gas measurement frequency. The rapid change in carbon dioxide production and oxygen consumption at the start and end of the feast-phase often falls between two measurements, this leads to an inherent uncertainty as shown in Figure 9B.

By allowing a larger process variance, the Particle Filter can respond to these fast changes, but will also more closely follow the off-gas measurements, resulting in a more jagged prediction of the OUR and CUR during periods with smooth transitions, as more prediction emphasis will come from the measurement variance. This is seen in the initial period of the famine phase, where some oscillations are seen in the carbon dioxide production rates. Determining the appropriate variances requires understanding of the biological process and measurement equipment. More frequent off-gas measurements assist the model as less variance builds up between two consecutive measurements.

#### Extending and adapting the model

Generally, the model predictions improve when additional independent measurements are available (42). One on-line measurement that is often available in aerobic bioreactors, but was not used in this model, is the dissolved oxygen (DO) concentration, which yields fast-responding continuous measurements, but suffers from signal drift, nonlinear probe dynamics, and loss in linearity due to fouling (43). de Jonge et al. (19) encountered DO related issues where, even in their experiments that lasted a couple of hours and involved rigorous calibrations, the DO probe data could not be used to improve the model performance. In the Supplementary Material 3 a variance smoothing algorithm is described which allows the high frequency, but possibly inaccurate, DO measurement to improve the predictions by utilizing the first order derivative of the dissolved oxygen measurement as virtual measurement. This measurement is used to scale the process variance, allowing the system to respond quickly and timely in periods with fast dynamics. This approach is only viable for aerobic systems, or for analogous probes that measure concentrations in the reactor broth. This addition to the Particle Filter demonstrates a significant benefit of this approach, compared to the more math heavy solutions of for example Farza et al. (12) and Oliveira et al. (13); i.e.: the relative ease by which the algorithm can be extended to new sensors, as well as adjusted to different bioreactor configurations and biological processes.

#### Initial state estimation

The initial state of the system is estimated by allowing *N* particles to assume uptake rates uniformly distributed around likely system states. By including more measurements before a system disturbance, the Particle Filter automatically converges towards the likely system state, and therefore the modeled uptake rates up to that point are prone to errors and should be discarded. The slowest convergence takes place for the hydrated carbon dioxide, especially at higher pH. Improved initial state predictions are produced by moving the start of state estimation backwards in time, if the dataset allows this, but it could also be derived mathematically by running the model backwards in time (not implemented here) (19). Other issues with the Particle Filter arise when additional processes take place beside biological. For example the exchange of a significant fraction of the broth with new medium can run into reconstruction issues, due to the unknowns of the new medium like temperature, pH, and the dissolved gas concentrations before it is added to the reactor. This is especially pronounced when reactor broth with a high carbonate concentration is exchanged for medium that is kept fully anaerobic by sparging it with nitrogen gas, and therefore has a low carbonate concentration. Wrongful or missing physical processes will translate in changes in off-gas measurements, which will erroneously be contributed to biological activity. Figure 9 right, shows this behavior to a minimal extent during the liquid exchange at t=11h. Generally, these events are easily spotted and corrected for by visual inspection of the reconstructed respiration rates.

#### Improved experimental design

The Particle Filter developed is a convenient tool for analyzing dynamically operated bioreactors and investigating the biological responses to the imposed dynamics, but it also allows optimizing the operational parameters of a cultivation to maximize the reconstruction resolution of the respiration rates. For example, by allowing the controlling system to adjust the gas flowrate of the cultivation it is possible to reconstruct respiration rates at high precision during phases with high and low respiration, without running into mass transfer limitations (examples are included in the Supplementary Material 5). The resulting dataset can then only be processed with a model as proposed here as an additional convolution is added to the off-gas signal.

Overall, the Particle Filter allows biotechnology researchers that focus on microbial cultivations to extend their research tools by facilitating reconstruction and interpretation of biological respiration rates with little effort. Researchers with specific and detailed interest in dynamic cultivations should be able to adapt the proposed model with kinetic models and additional on-line analysis that are available to them.

## Conclusions

A generalized respiration rate reconstruction tool is developed for dynamic operated bioreactors, based on the Particle Filter method. The reconstructed respiration rates are a direct reflection of biological activity in the bioreactor, and can be used to characterize the system developments over time. As a specific example, the functional characterization of a pulse fed bioreactor throughout an enrichment of 100 days was achieved solely based on on-line data. This allowed for the identification of shifts in the dominant metabolic processes without prior knowledge or additional off-line measurements of the system. Furthermore, the general model was able to reconstruct the respiration rates in many differently operated sequencing batch bioreactors. This enables the identification of specific process properties that define the development in time of the process. This paves the way for future implementations where the development in time of the microbial community structure and the functional performance can be linked at high resolution providing crucial insights in the factors that shape the microbial world

## ACKNOWLEDGEMENTS

We gratefully acknowledge financial support from the Netherlands Organization for Scientific Research (NWO) funded UNLOCK project (NRGWI.obrug.2018.005) and the company Paques B.V. (Paques Partnership Program project 13002).

## Supplementary Material 1: Parallel bioreactor setup

**Fig. S1.**
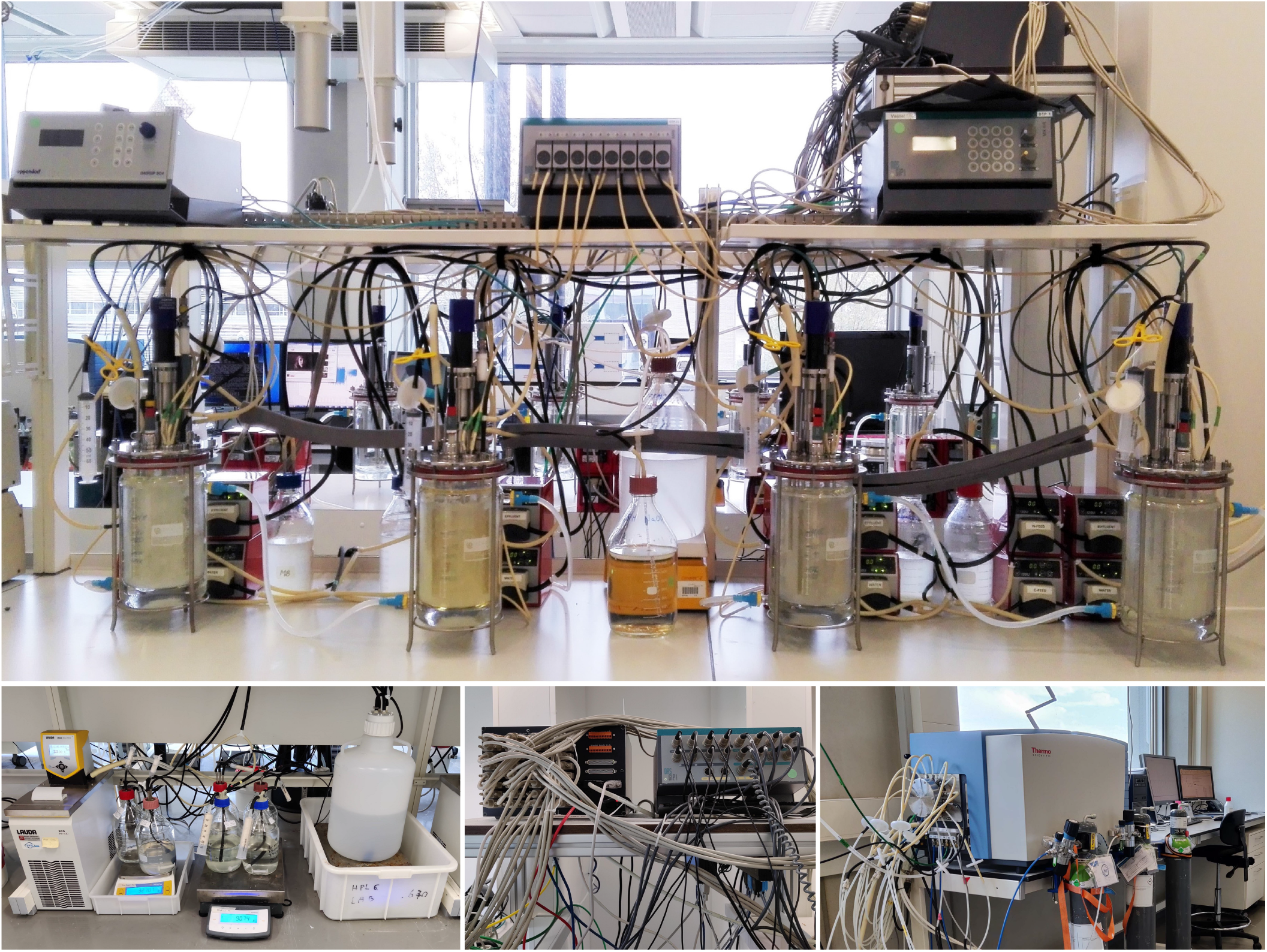
Photos of the parallel bioreactor setup. Shown are (top) four of the eight bioreactors, with their (bottom) balances and cultivation media, the custom central control system that connects all digital and analog equipment, and the off-gas mass-spectrometer with channel selector and calibration gasses.

## Supplementary Material 2: Particle Filter implementation

The implementation of the Particle Filter, physicochemical model, and data pruning are included in the on-line code repository at: https://github.com/GRS-TUD/d2i, including data examples. What follows is a construction of the most important steps of the general implementation, which are useful for custom implementations and extensions. The reconstruction of the respiration rates occurs through the use of a Particle Filter. The idea behind the Particle Filter is to perform a set of Monte Carlo simulations in parallel, and then to perform Bayesian inference on the predicted system state and the available measurements. If the number of particles is large, and the model that describes the system is accurate, it becomes possible to identify hidden (non-measured / non-measurable) states. In most microbial cultivations one of those hidden states is the respiration rate of the microorganisms. The available measurement is relatively far removed from this state in the form of off-gas measurements (see section Physicochemical processes for gas exchange).

For example: oxygen is taken up by microorganism from the liquid broth at a specific rate 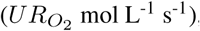, this causes a decrease in dissolved oxygen 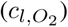. The rate of gas exchange between the bubbles and the liquid depends on gas transfer coefficient 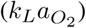 and the difference of concentration in the liquid and partial pressure of oxygen in the bubbles 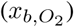, corrected for the Henry coefficient 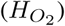. The concentration in the gas bubbles depends on the inflow gas rate (*F*_*in*_, dry air), the head space recycling gas rate (*F*_*rec*_), and their relative compositions (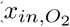 and 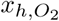). The exchange of gas molecules between the bubbles and the liquid changes the molar gas flow from bubbles going in (*F*_*in*_ + *F*_*rec*_) and coming out of the liquid (*F*_*b*2*h*_). The gas coming from the bubbles mixes with the gas in the head space. The system is kept at operation pressure (*P*), which requires a net gas outflow from the reactor to the off-gas measurement equipment. To that end the gas is dried (*P*_*w*_), and the gas composition is measured.

Comparable processes occur for all gaseous compounds that are consumed or produced in a bioreactor. All gasses affect the relative concentration of the other gasses, therefore the reconstruction needs to take all gasses into account - including gasses that do no partake in biological processes in the system. Two additional significant complications arise due to carbon dioxide speciation, where a significant fraction of CO_2_ is dissolved in the liquid as carbonates. The second complication is caused by system dynamics, where either physical processes, like liquid exchange, environmental change (pH, temperature, pressure), gas change, or biological processes, like changing metabolisms, response to disturbances, or succession are causing rapid and slow changes in the measurements. If all physicochemical processes are taken into account, then all changes that remain can be ascribed to changes in biological rates. And such is the goal of the Particle Filter with physicochemical model described in this work. Below follows the implementation of the Particle Filter, where the physicochemical aspect is covered in the main text.

**(Particles)** Let P = [*p*_*ij*_] be a matrix of size v*×* N (i.e., N represents the number of particles and v represents the number of states of the system and is equal to (4 *·w* + 1), with w the number of gas compounds of interest). For each gas the states include an uptake rate, concentration in the liquid, fraction in the bubbles, and fraction in the head space (i.e., total of 4 states per gas compound). One additional system state is required to describe all hydrated carbon dioxide species.

**(Initial state)** Let D = [*d*_*pq*_] be a matrix of size w *×* M (i.e., w represents the number of gas compounds measured by the off-gas analysis, and M the number of independent measurements. The initial state estimate assumes equilibria between the head space, bubbles, and liquid phase (equilibrium calculations based on environmental conditions and Henry coefficients), this populates P from *p*_*iw*_ onwards. All particles can start with identical state estimates, or an exception can be made for the uptake rates. The initial uptake rates *p*_*i*0_ till *p*_*iw*_, then need to be picked uniformly between a ceiling and floor vectors of size w. This process improves the calculation time until a proper estimate of the uptake rates is achieved. The time *t*_*k*_ is set to the time of the initial measurement *t*_0_.

**(Uptake Rate covariance)** Let U be a matrix of size w *×* w (i.e., often a diagonal matrix of the rate by which the uptake rates can change per time step). This covariance matrix determines the average variation that occurs each time step for the uptake rates. When elements are set to 0, the uptake rates will remain constant throughout the modeling, larger values allow for more disperse particles. If relationships exist between gas compounds (e.g. oxygen and carbon dioxide in heterotrophic systems, they can be included in de covariance matrix as it restrict the creation of unlikely system states). The determination of the covariance matrix values requires feeling for the system that is modelled.

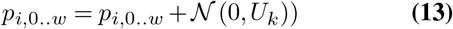

The uptake rates in the particle filter states are updated after each calculation step with a normally distributed random value based on the uptake rate covariance matrix U. The values of the covariance matrix can also be made dependent on time (*k*), as demonstrated in Supplementary Material 3. Additional checks can be included that limit the range of the uptake rates, e.g.: only positive or negative uptake rates for specific gas compounds. But particles that show biologically impossible behavior will easily be removed in the resampling step. Therefore no specific knowledge of the biological conversions is required at the expense of additional computational calculations.

**(Predict)** For each measurement m from 0…M we retrieve the time stamp and calculate the number of prediction cycles that have to be completed before a new measurement is available (i.e., from *t*_*k*_ till *t*_*n*_). During each prediction step (of 1 second) one full iteration of the physicochemical model is made on P, which yields predictions for each particle, based on its current system state. After each model step, the uptake rate states in P are allowed to vary according to the Uptake Rate covariance matrix. This process is repeated until the next measurement is available, if measurements are sparse the predictions of the particles will vary significantly.

**(Measurement covariance)** Let R be a matrix of size w*×* w (i.e., a diagonal matrix of the variance of the standard error of the measurement equipment). This matrix is relatively easy to construct but might be subject to change if the equipment results in different errors depending on the relative or absolute nature of the error. Most off-gas analyzers cannot measure well at both low and high percentages of the same gas, given the same calibration. Any knowledge on expected measurement variance is included in the model through the measurement covariance matrix R.

**(Update)** After all predictions between two measurements have been completed, the predicted state estimates are compared to the real measurements. The state estimates contain a prediction for the head space gas fraction, predictions that are close to the measured values are more likely to be correct. The distance between the measured and predicted values is expressed as the likelihood that the predication occurred given the uncertainty of the measurement covariance matrix R. To this end the probability density function (PDF) for a normal continuous random variable is used.

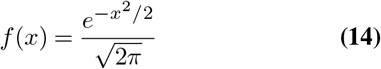

If many prediction steps exist between to measurement, and multiple gas compounds need to be modeled, it is likely that a large fraction or all of the predictions are very far removed from the measurements. From a computational point of view we therefore make use of the logarithmic PDF, as it allows for a wider coverage in uncertain areas (300 orders of magnitude). Each particle has a weight attributed to it (*W* of size N), which represents how likely this particle is to be the real representation of system state. The weights are updated with the new PDF and the total is normalized to 1.

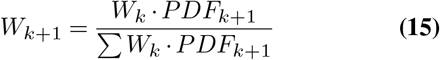

**(Estimate)** The updated weights are used to make an estimate of the current system state by taking a weighted average of the particles. Also the weighted variance of the estimate can be calculated in an analogous fashion. These estimates represent the mathematically best prediction of the system states.

**(Resample)** Importantly, an ever increasing fraction of the particles is assuming system states that are far removed from the likely states. Resampling is required to stop needless calculations and to improve next predictions by utilizing the unlikely particles as copies of more likely particles. The resampling threshold and resampling method allow for control on this process. Here we chose to implement the systematic resampling algorithm which yields new particles proportional to their weight. After the resampling step several copies of a particle exist, but after the prediction step they are all updated with a different variance in their uptake rate estimates, which allows for a finer scanning around the current best system state estimates.

**(Environmental triggers)** The physicochemical model needs to take into account all imposed changes that occur throughout a cycle, as they are likely to affect the measurements. If these imposed changes are not modelled well they will skew the reconstruction of the respiration rates. To this end, several (time) triggers can be included in the model that allow for changes in environmental parameters (e.g., reactor volumes, liquid exchange (including dissolved gasses), gas flows, in-gas composition, ionic strength). Parameters depending on temperature, pressure, and ionic strength are automatically adjusted (e.g.: Henry coefficients, kLa, and activity coefficients). These time triggers can also be used to dynamically change the Uptake Rate covariance matrix U, or measurement covariance matrix R, based on system specific knowledge.

## Supplementary Material 3: Adding DO measurement to Particle Filter

A straightforward implementation of the DO probe in the Particle Filter would be to use it as measurement of 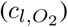. The DO measurement is available at each prediction step, and could therefore yield significant improvements to the estimates of the particles, as they can be steered over 100 times more often than only based on off-gas measurements. The downside of the DO probe is that it is susceptible to signal drift and loss in linearity, especially in longer cultivations and under dynamic conditions. Therefore, if the DO measurements are included, even with significant large error margins in the measurement covariance matrix R, it will have a tendency to pull the particles towards its erroneous representation of 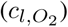. This behavior is observed in state estimation filters when precise measurements, with variable accuracy, are included.

Alternatively the precision and frequency of the DO measurements can be used as environmental input to dictate the Uptake Rate covariance matrix U for each time point. In periods where the DO measurements do not change (dDO/dt≈ 0), it is also more likely that the respiration rates change little. And during periods with strong changes in DO, the direction of the change can be used to indicate an proportional increase or decrease in oxygen respiration rates. By combining the Ramer-Douglas-Peucker algorithm (Supplementary Material 4), or a general signal smoothing algorithm, the DO data can be used to improve the respiration rate reconstructions to dynamics occurring at 5 to 10 second resolution. Investigations where cultivations are exposed to stronger dynamics are required to verify the reconstructed rates.

The relationship between U_k_ and the rate of change in the dissolved oxygen concentration is mechanistically linear. Yet, with respect to the Particle Filter it is always preferable to allow some variance in the uptake rate kinetics.

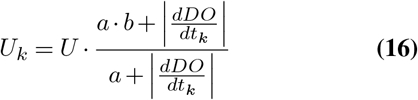

Saturation curve with half saturation parameter *a* and offset parameter *b*, which prevents U_k_ from becoming 0.

## Supplementary Material 4: Determination of the feast-length

We determined the end of the feast phase based on the inflection points of the OUR, CER, and the on-line measured dissolved oxygen (DO) profile. In several rapid sampling experiments we measured the acetate profiles of different reactors. Interestingly, the DO plots, and off-gas profiles during intense sampling look significantly different from undisturbed cycles. This is likely caused by the interference from sampling, and the decreasing reactor volume.

**Fig. S2.**
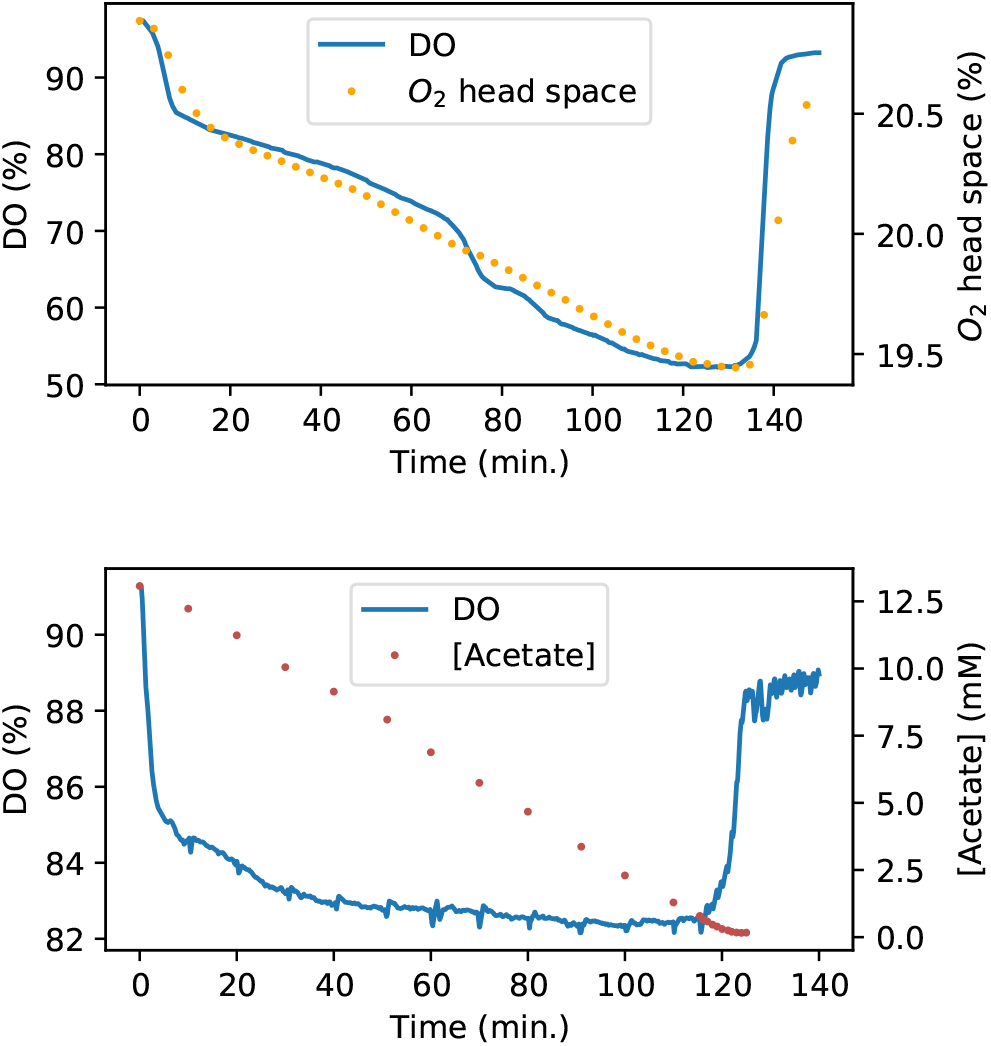
DO and *O*_2_ concentration in the head space of an undisturbed feast famine reactor (top) after a pulse at t=0, and the acetate and DO profiles during the following operation cycle (bottom). Difference in DO profiles between the two cycles likely results from sampling, and decreasing reactor volume, as the following (undisturbed) cycle more closely represents the top figure (not shown).

Figure S2 top, shows a typical oxygen and DO profile associated with a growing culture. A clear deflection point is visible around t=140 where the respiration peaks and quickly decreases. This point is correlated with low extracellular acetate concentrations (bottom), followed by complete consumption of acetate. Comparable graphs and data were collected for reactors that exhibited the typical hoarding strategy, and of cultures exhibiting partial growth and hoarding characteristics. From these measurements we could derive and verify a mathematical definition for the feast phase length.

**Fig. S3.**
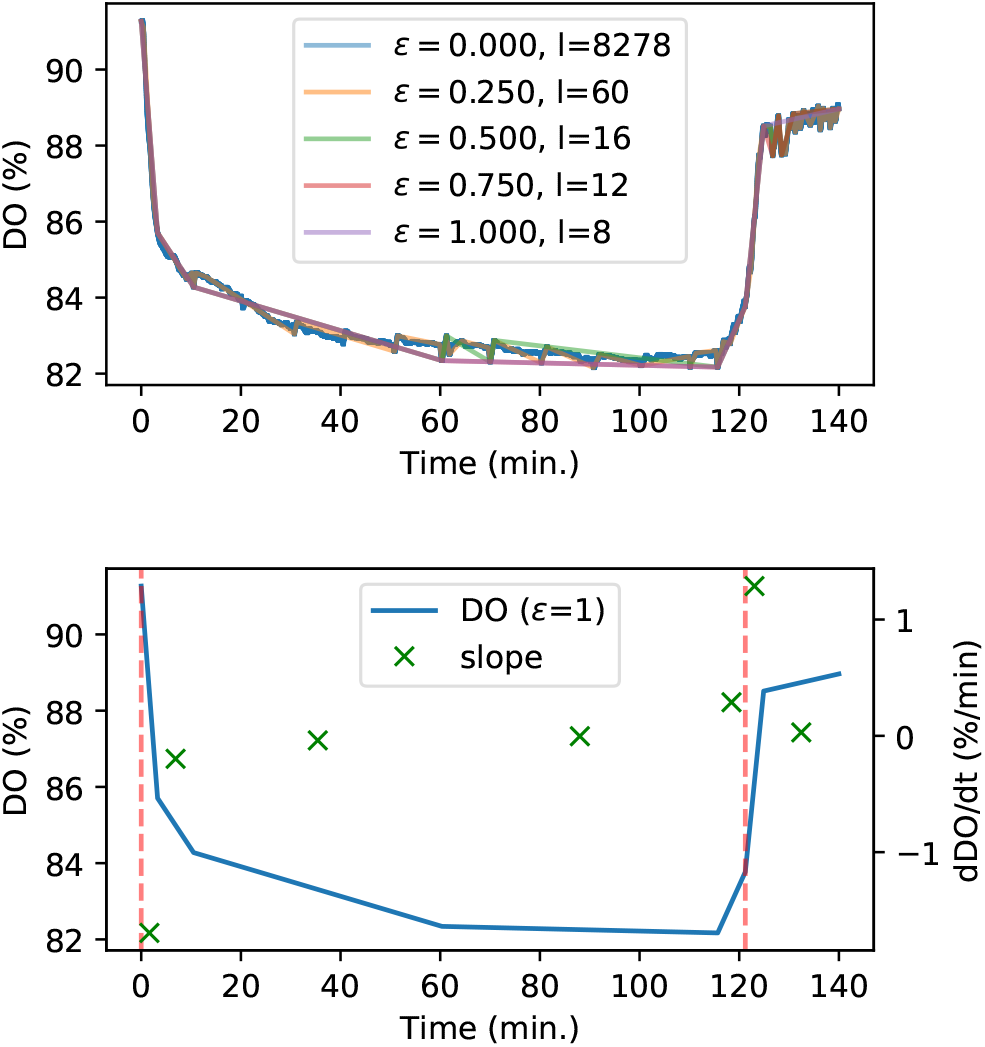
Identification of significant time points from the original DO dataset with the Ramer-Douglas-Peucker algorithm. It allows for a close estimate of the beginning and end of the feast phase (dashed red lines) by selecting the fragments with the minimum and maximum slope (green x).

The DO dataset contains a measurement every second, these 8400 data points (140 min.) together describe the DO curve in Figure S2 (bottom). The Ramer-Douglas-Peucker algorithm (36) can be used to reduce the number of points while still capturing the shape of the curve. The algorithm requires one fitting parameter ϵ which is the maximum distance any data point can have from the approximated curve. In Figure S3 top, the overlapping RDP curves are plotted with 5 different values for ϵ. In this manner, it becomes possible to identify important points; the original curve can for example be approximated by 8 data points, given that all other points are within 1 DO %. And alternatively, an ϵ of 0.25 allows for easy identification of all sampling points. In either case, the end of the feast phase most closely correlates with the section with the steepest slope (Figure S3 bottom).

Analogue calculations can be made for the reconstructed OUR and CER, which are based on less frequent, but independent, off-gas measurements. And in most dynamic systems, the same approach can be used for acid or base titration data, or gradual shifts in pH within the dead band of pH controlled reactors.

## Supplementary Material 5: Increase off-gas measurement resolution

To improve the reconstruction accuracy a significant change in gas composition of in-gas and off-gas is beneficial. A lower gas inflow rate results in a stronger change, but also comes at the cost of a stronger delay and convolution of respiration dynamics. Additionally, a different gas inflow rate also changes the partial pressures of the gasses, which can lead to significant changes in the speciation of compounds, and even to limitations. The conflicting requirements regarding improved off-gas signal and capturing of process dynamics, preferably without disturbing the system must be considered during experimental design.

By allowing for changing gas inflow rates, the resulting off-gas signal becomes exceptionally more difficult to interpret without a physicochemical model. The proposed model in this work allows for experimental design with changing flow rates, and changing in-gas compositions. In Figure S4 an example is shown where during the famine phase of a feast-famine system little respiration activity is taking place compared to during the feast phase. This results in a strong limitation on the ability to reconstruct the biological respiration rates as the off-gas signal comes within the measurement uncertainty of the equipment. By reducing the gas inflow rate instantaneous after 5 hours of cultivation, the signal is ‘amplified’ and reconstruction is possible with higher fidelity. Naturally, more complex and more autonomous changes are possible regarding the in-gas flow and composition.

**Fig. S4.**
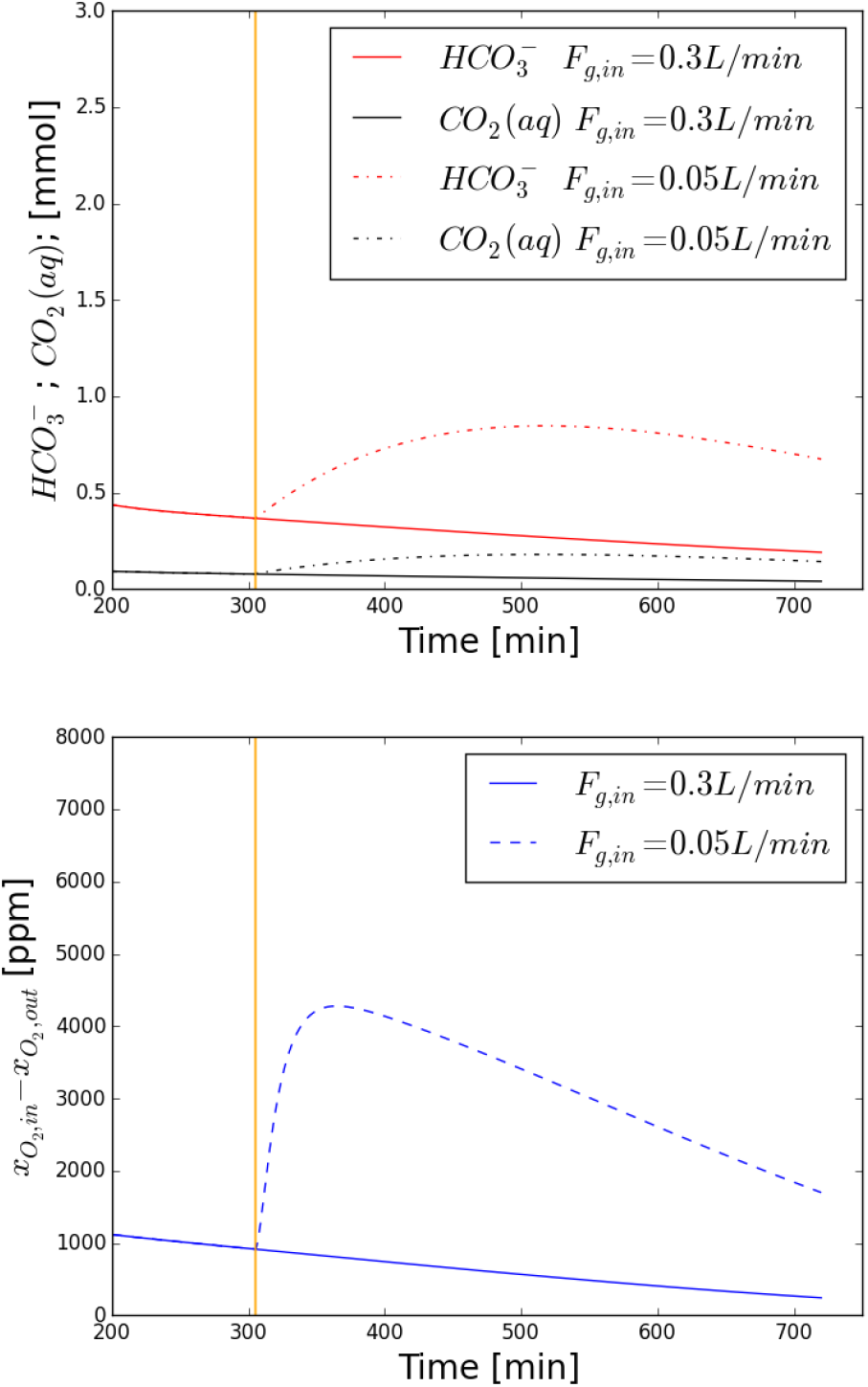
Simulation of the liquid *CO*_2_ and 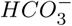, and the gas phase *O*_2_ profiles; with constant and adapted gas inflow rates. The orange line indicates where the flow rate is changed from 300 to 50 mL min^-1^.

## Footnotes

1 Note that Wang and colleagues swapped the naming convention for the equilibrium constants Ka1, and Ka2, and inadvertently allowed some printing errors in the Parameters section of the Materials and Methods regarding the relationships between the equilibria.

